# Levodopa Responsiveness Subtypes of Freezing of Gait: Results Using a Levodopa Challenge

**DOI:** 10.1101/667071

**Authors:** J. Lucas McKay, Felicia C. Goldstein, Barbara Sommerfeld, Douglas Bernhard, Sahyli Perez Parra, Stewart A. Factor

## Abstract

**Objective:** To demonstrate that levodopa-unresponsive freezing of gait (ONOFF-FOG), which is a disabling untreatable feature of Parkinson’s disease (PD), is distinct from responsive/OFF only FOG (OFF-FOG) and potentially has a distinct pathophysiology.

**Methods:** Fifty-five PD patients completed levodopa challenges after >12 hours OFF with supratherapeutic doses to classify them as NOFOG, OFF-FOG or ONOFF-FOG. Serum levodopa levels ensured threshold levels were met. An “ON” response was defined as ≥20% improvement in MDS-UPDRS-III score. Main outcome measure was MDS-UPDRS-III based on clinical exam, timed-up-and-go tests and 360° turns.

**Results:** 45 patients exhibited an “ON” response to the challenge. Levodopa-equivalent-dose was 142±56% of patients’ typical morning doses. Patients could be classified as: 19 ONOFF-FOG, 11 OFF-FOG, 15 NOFOG. The ONOFF-FOG group exhibited significantly higher NFOG-Q, MDS-UPDRS-II/III scores compared to the OFF-FOG group. MDS-UPDRS-III total varied significantly across medication response states (P<0.01) and groups (P=0.03). Among MDS-UPDRS-III subdomains, significant effects of group (highest in ONOFF-FOG group) were identified for measures of lower extremities and gait. Linear mixed models revealed a highly significant association between serum levodopa level and MDS-UPDRS-III score that did not vary across the groups. There was a highly significant group interaction effect on the association between serum levodopa level with MDS-UPDRS-III item 11 “Freezing of Gait” (P<0.001).

**Conclusions:** ONOFF-FOG is a distinct subtype of FOG in PD, as opposed to OFF-FOG. This is an important step in demonstrating that FOG is not a single entity and, in turn, could lead to better understanding of pathophysiology and development of effective therapies.

## Introduction

Freezing of gait (FOG), described as a brief arrest of stepping when initiating gait, turning, and walking straight ahead, is a common, poorly understood symptom complex that has potentially grave consequences to Parkinson’s disease (PD) patients^1, 2^. It is unpredictable in character, is a leading cause of falls and consequent injuries, results in loss of independence and social isolation, and treatment thus far is limited^1, 3^. There has been great variability in findings related to physiological, imaging, motor and non-motor correlates as well as therapeutic response to various treatment modalities^4^. FOG appears to develop and/or progress independently of the other motor features, is associated with specific risk factors, and is thought to be caused by specific but as yet unknown pathology^5^. Recognition that FOG is not a single uniform symptom but instead exists in several subtypes is crucial to improved understanding of the pathophysiology and development of new therapies^6^.

Clinical subtypes have been the subject of discussion. The phenomenology of FOG differs between patients, and may include shuffling with small steps, trembling in place without forward movement, or total akinesia^7^. In addition, the settings such as starting, turning or walking and the effect of special constraints (such as going through doorways) also differs between patients. It remains an open question as to whether these phenomenologies are the result of different severity or represent different pathophysiologies.

One other possible clue to potential classification of FOG may relate to levodopa responsiveness. The relationship between FOG and levodopa response is complex. It is suggested that several apparent subtypes exist including: 1) FOG which appears only in the “OFF” state, and disappears in the levodopa induced “ON” state (referred to here as OFF-FOG); 2) FOG that is unresponsive and is present in the “OFF” and “ON” state (ONOFF-FOG); and 3) FOG present during “ON” state and absent in the “OFF” state (drug-induced or ON-FOG)^8 7 9 10^. One study indicated that 62% of FOG patients are OFF-FOG, 36% ONOFF-FOG and 2% ON-FOG^11^.

It has been questioned whether these subtypes, particularly ONOFF-FOG, are distinct pathophysiologically or if the “on” FOG in those with ONOFF-FOG represents undertreatment^7,9,11^. As this is unresolved, some also suggest that FOG progresses over time from responsive OFF-FOG to unresponsive ONOFF-FOG, indicating that subtypes do not actually exist^3, 4^ Others, on the other hand, have shown that ONOFF-FOG can come on without prior OFF-FOG^9^ suggesting they may have different physiology^4^. What is certain is that the relationship of FOG to levodopa therapy has been inadequately examined to this point.

For the purpose of this study, we hypothesize that separate levodopa responsive subtypes of FOG exist and that ONOFF-FOG is a distinct form that is not the result of inadequate therapy. We also intend to demonstrate that FOG of the ONOFF-FOG type differs from other PD cardinal features pharmacologically in that it has limited responsiveness. In order to concretely establish that ONOFF-FOG is truly a symptom distinct from OFF-FOG and that ONOFF-FOG behaves differently from other parkinsonian symptoms in response to levodopa, we examined the nature of levodopa responsiveness of PD patients with and without FOG using a levodopa challenge paradigm. We examined patients in the practically defined “OFF” state and after a levodopa equivalent dose greater than their typical morning dose of medications. We measured serum levodopa levels to establish that they reached adequate threshold levels indicating that any remaining FOG was not a consequence of inadequate levodopa dosage or delayed efficacy due to poor gut absorption. Our expected outcome was that patients with ONOFF-FOG would exhibit responses in overall parkinsonian symptoms, as assessed with MDS-UPDRS-III^12^, to changes in serum levodopa level that were comparable to those observed in patients without FOG (NOFOG) and with OFF-FOG. We also comprehensively investigated whether the response of individual parkinsonian symptoms to medication state varied across FOG groups.

## Methods

### Study population

All participants were recruited from the Emory Movement Disorders clinic and provided written informed consent according to procedures approved by Emory University IRB. The study population included PD patients with and without FOG. Inclusion criteria for all participants were: Age≥18 years; PD diagnosis according to United Kingdom Brain Bank criteria^13^; Hoehn & Yahr stage I-IV in the OFF state; Demonstrated response to levodopa; Able to sign a consent document and willing to participate in all aspects of the study. Additional inclusion criteria for participants with FOG were: FOG noted in medical history and confirmed visually by examiner in the office. Exclusion criteria for all participants were: Atypical parkinsonism; prior treatment with medications that cause parkinsonism; neurological or orthopedic disorders interfering with gait; dementia or other medical problems precluding completion of study protocol.

### Clinical and demographic variables

Clinical and demographic data were collected using a battery of standardized instruments. PD duration, FOG duration and ages at PD and FOG onset were taken via self-report. Self-reported FOG severity was assessed with the New FOG Questionnaire (NFOG-Q), and patients completed a falls diary. Full MDS-UPDRS were also completed.

### Levodopa challenge paradigm

All patients came to clinic in the practically defined “OFF” state >12 hours after last intake of antiparkinsonian medications. They were assessed for motor symptoms using the MDS-UPDRS-III and three timed-up-and-go tests; standard, cognitive and carrying a tray with cups on it. They also completed two 360° turns in each direction. These were completed in the motion capture lab and videoed. After assessment completion all patients were administered a levodopa equivalent dose (LED) greater than their typical morning dose. Dosage amounts were individualized to each patient by the examining investigator (SAF) based on whether dyskinesia was an issue and the size of their typical dose. For example, if a patient had a known moderate dyskinetic response to their usual dose, a modest increased dose was given. If they had no or minor dyskinesia they were given a dose up to 200% of their usual dose. Patients were subsequently assessed at regular intervals until they reached their full “ON” at which point the “ON” examination was completed (same measures as done in the “OFF” state). The interval between “OFF” and “ON” testing varied from 30 minutes to 3 hours. Blood was drawn for measurement of levodopa level during the “OFF” state and immediately preceding the full “ON” assessment.

In order to increase the likelihood of producing FOG^14^, the gait assessment of the MDS-UPDRS-III was amended to include the common clinical protocol in which the patient walks in a straight line, turns around and returns to the examiner, as well as with the following additional conditions: timed-up-and go^15^ with and without a dual task^16^, and rapid 360° turns^14^. Performance was scored in-person and scores were confirmed from video and amended if necessary (12/110 total scores).

We classified the response of each patient to the levodopa challenge as clinically-meaningful or not based on an observed improvement of ≥20% in MDS-UPDRS-III score after levodopa administration^17^. Plots of improvement vs. MDS-UPDRS-III OFF score were examined for potential associations between improvement and baseline score or study group.

### Levodopa levels

A previously reported method was followed^18^. HPLC with ESA 5600A CoulArray electrochemical detection system, equipped with an ESA Model 584 pump and an ESA 542 refrigerated autosampler was used. Separations were performed at 35°C using an MD-150 × 3.2 mm C18 column equipped with a C18 column guard cartridge.

### FOG group assignment

We classified each patient into one of three study groups determined *a priori* based on history of the presence of FOG along with scores on the MDS UPDRS-III item 11 “Freezing of Gait” exam in each of the “OFF” and “ON” medication states. Participants who had no history of FOG and received a score of zero on item 11 in both medication states were classified as “no freezing,” or “NOFOG.” Participants who received a nonzero score on item 11 in the “OFF” medication state but a zero score in the “ON” medication state were classified as “OFF-FOG.” Participants who received a nonzero score on item 11 in both medication states were classified as “ONOFF-FOG.” All patients were categorized into study groups, independent of whether they exhibited a clinically-meaningful response to the levodopa challenge test. Only patients that exhibited a clinically-meaningful response to the levodopa challenge were included in cross-group analyses.

## Statistical analysis

### Demographic and clinical characteristics

We assessed differences in clinical and demographic characteristics across all three study groups with ANOVA and Tukey tests or chi-squared tests as appropriate. Comparisons of demographic and clinical variables specific to FOG (age at FOG onset, FOG duration, NFOG-Q score) were performed only among the two FOG groups.

### Motor features and serum levodopa levels across groups in the “OFF” and “ON” states

We used two approaches to analyze changes in parkinsonian symptom level (MDS-UPDRS-III total score and subdomains) and serum levodopa level between the “OFF” and “ON” medication state in each group. First, we used separate repeated measures ANOVAs to compare MDS-UPDRS-III scores and serum levodopa levels in each of the “OFF” and “ON” medication states across groups, with PD duration included as a covariate to control for imbalances across groups. MDS-UPDRS-III total scores were calculated with and without item III.11, “Freezing of Gait” as this item was used to establish group allocation. In these analyses, items with multiple subscores (e.g., item 16, kinetic tremor of the left and right upper limbs) were assembled into total subscores prior to analysis.

Next, we used linear mixed models in SAS PROC MIXED to determine whether associations between MDS-UPDRS-III scores and serum levodopa level differed across the NOFOG, OFF-FOG, or ONOFF-FOG groups. In these analyses, PD duration was included as a covariate and separate random intercepts and slopes were calculated for each patient. The OFF-FOG group was specified as the reference group to enable contrasts between the OFF-FOG and ONOFF-FOG groups. Satterthwaite’s approximation was used to estimate denominator degrees of freedom as necessary. Linear combinations of model coefficients were estimated with approximate *t*-tests. Statistical tests were performed at alpha ≤ 0.05 in SAS University Edition 7.2.

## Results

### Levodopa challenge test

Of N=55 patients enrolled, N=45 (82%) exhibited a clinically-meaningful response to the levodopa challenge (≥ 20% decrease MDS-UPDRS-III; Figure 1A). The average improvement in total MDS-UPDRS-III score was 47±15% (range, 20–80%) among patients who exhibited a clinically-meaningful response and 3±14% (−23–19%) among patients who did not. No clear associations between improvement in MDS-UPDRS-III score or study group and baseline score were apparent on inspection of plots of improvement vs. OFF score (Figure 1B). The average administered LED was 395±243 mg (range, 133-1348 mg) in the levodopa challenge, corresponding to 142±56% (range, 62-325%) of patients’ typical morning dose. Administered doses did not significantly differ among those patients who did or did not exhibit a clinically-meaningful response (levodopa-equivalent-dose in mg, P=0.81; LED as proportion of morning dose, P=0.29). Serum levodopa levels after levodopa administration were significantly higher (79%, P<0.01) among patients who did not exhibit a clinically-meaningful response (Table S1).

**Figure 1.**
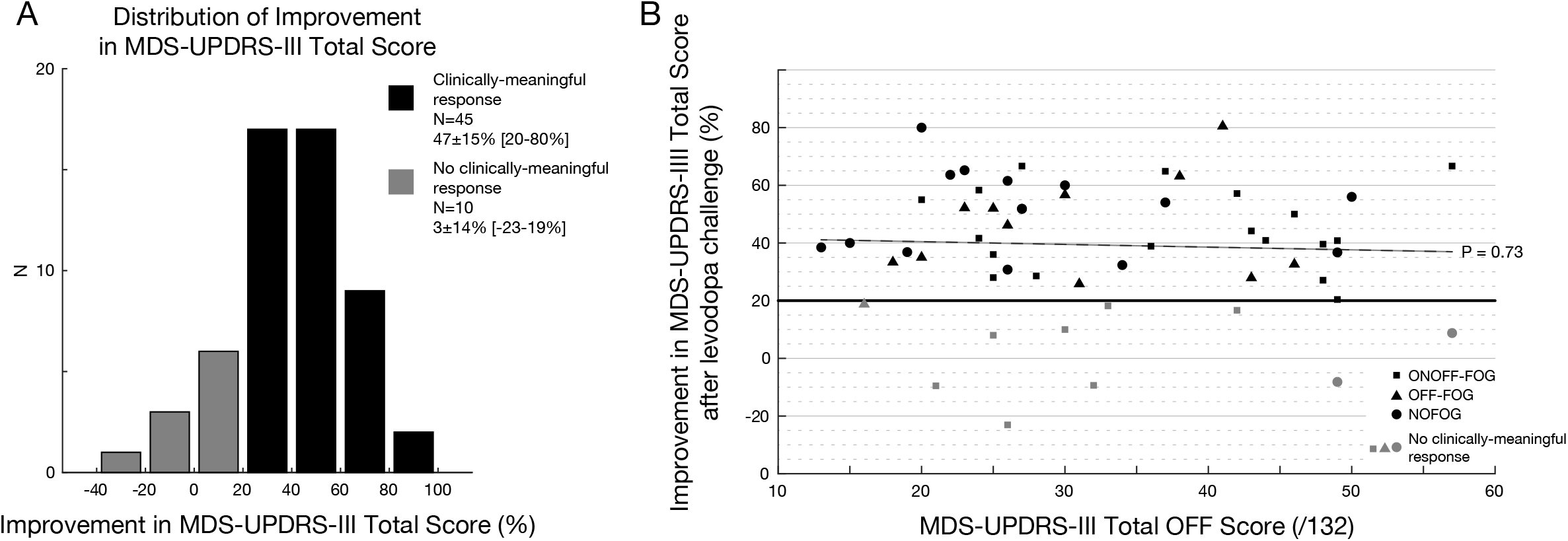
Improvement in MDS-UPDRS-III total scores after acute levodopa challenge. A: Distribution of reduction in MDS-UPDRS-III total scores among patients who did and who did not exhibit a clinically-meaningful response (black and gray bars, respectively). B: Scatterplot depicting association between reduction in MDS-UPDRS-III total score after levodopa challenge and OFF medication score. Colors as in A. Best fit trendline for entire sample is shown for reference (dashed).

### Demographic and clinical characteristics of 3 groups

Demographic and clinical characteristics of patients who exhibited a clinically-meaningful response to the levodopa challenge are summarized in Table 1. Of N=45 patients, 19 (42%) were in the ONOFF-FOG group, followed by NOFOG 15 (33%), and OFF-FOG 11 (25%). None of the patients exhibited levodopa-induced ON-FOG^9^. No significant differences between groups were observed in age, sex, education, MoCA score, age at PD onset or FOG onset, FOG duration, presence of dyskinesia during the “ON” state (as indicated by the investigator while completing the MDS-UPDRS-III) or MDS-UPDRS-I or IV subscores.

**Table 1.** Demographic and clinical characteristics of the study sample, overall and stratified on FOG status.

| Characteristic | All<br>N=45 | NOFOG<br>N=15 | OFF-FOG<br>N=11 | ONOFF-FOG<br>N=19 |
| --- | --- | --- | --- | --- |
| Age (y) | 66±8 | 65±11 | 68±4 | 67±7 |
| Sex |  |  |  |  |
| Male (n,%) | 33 (73) | 9 (60) | 9 (82) | 15 (79) |
| Female (n,%) | 12 (27) | 6 (40) | 2 (18) | 4 (21) |
| Education (y) | 16±2 | 17±1 | 17±3 | 16±2 |
| MoCA (/30) | 25.4±4.0<sup>a</sup> | 27.4±1.9 | 25.1±4.2 | 24.0±4.7 |
| PD duration (y)* | 10±6 | 6±4<sup>†</sup> | 11±5 | 12±7<sup>†</sup> |
| Age at PD onset (y) | 56±9 | 58±11 | 57±6 | 55±9 |
| LED (mg)** | 1294±664 | 864±300<sup>†</sup> | 1197±500 | 1690±738<sup>†</sup> |
| MDS-UPDRS-I (/52) | 11.8±6.3 | 10.2±5.1 | 11.5±5.5 | 13.2±7.5 |
| MDS-UPDRS-II (/52)* | 16.0±7.7 | 10.7±6.9<sup>†</sup> | 17.3±5.7 | 19.3±7.3<sup>†</sup> |
| MDS-UPDRS-IV (/24) | 6.3±3.2 | 5.5±3.1 | 6.3±3.4 | 7.1±3.3 |
| ON-state dyskinesia |  |  |  |  |
| Yes (n, %) | 34 (76) | 9 (60) | 10 (91) | 15 (79) |
| No (n, %) | 11 (24) | 6 (40) | 1 (9) | 4 (21) |
| FOG duration (y) | 3±3<sup>b</sup> |  | 4±3 | 3±3 |
| Age at FOG onset (y) | 64±7<sup>b</sup> |  | 64±6 | 64±7 |
| NFOG-Q (/28)** | 20.1±5.5<sup>b</sup> |  | 16.5±6.1<sup>†</sup> | 22.1±4.0<sup>†</sup> |

Abbreviations: NOFOG, no Freezing of Gait; OFF-FOG, levodopa-responsive Freezing of Gait; ONOFF-FOG, levodopa-unresponsive Freezing of Gait. \*Overall effect of group at P<0.05. \*\*Overall effect of group, P<0.01. <sup>†</sup>Significant difference between listed groups at P<0.05. <sup>a</sup>N=43. <sup>b</sup>N=30.

Significant contrasts were observed between the ONOFF-FOG and NOFOG groups on PD duration (12±7 vs. 6±4 y, P<0.01) and daily LED (1690±738 vs. 864±300 mg, P<0.01). No significant differences were observed between the OFF-FOG and ONOFF-FOG groups on these variables. The ONOFF-FOG group exhibited significantly more impairment on NFOG-Q (P<0.01) and MDS-UPDRS-II (Motor Aspects of Experiences of Daily Living) (P<0.01) compared to the OFF-FOG group. Demographic and clinical characteristics of all patients enrolled, whether or not they exhibited a clinically-meaningful response to the levodopa challenge, were very similar overall and are summarized in Table S2.

Levodopa doses administered during the levodopa challenge did not vary across groups (P=0.50) (Table 2). When calculated as proportion of the typical morning dose, administered doses were higher in the OFF-FOG group compared to the ONOFF-FOG group (181% vs. 121%, P=0.02). Doses were also higher compared to the NOFOG group; however, this difference was not statistically significant.

**Table 2.** Levodopa responsiveness of the study sample, overall and stratified on FOG status.

| Characteristic | All<br>N=45 | NOFOG<br>N=15 | OFF-FOG<br>N=11 | ONOFF-FOG<br>N=19 |
| --- | --- | --- | --- | --- |
| Levodopa challenge dose |  |  |  |  |
| Levodopa equivalent (mg) | 391±260 | 282±133 | 438±349 | 451±263 |
| Proportion of morning dose (%)<sup>*</sup> | 138±59 | 119±31 | 181±92<sup>a</sup> | 128±38<sup>a</sup> |
| Serum levodopa<sup>¶,†,‡</sup> |  |  |  |  |
| OFF (ng/mg) | 0.3±0.4 | 0.1±0.1 | 0.4±0.3 | 0.5±0.5 |
| ON (ng/mg) | 27.9±16.8 | 19.9±11.3<sup>a</sup> | 28.0±11.9 | 34.8±20.8<sup>a</sup> |
| MDS-UPDRS-III<sup>*,†</sup> |  |  |  |  |
| OFF (/132) | 32.5±11.3 | 27.9±10.9<sup>a</sup> | 31±9.7 | 36.9±11.2<sup>a</sup> |
| ON (/132) | 17.2±8.3 | 13.9±7.2 | 16.5±8 | 20.3±8.6 |
| MDS-UPDRS-III (item 11 excluded)<sup>†</sup> |  |  |  |  |
| OFF (/128) | 30.8±10.7 | 27.9±10.9 | 29.4±9.7 | 34.1±10.8 |
| ON (/128) | 16.6±8.1 | 13.9±7.2 | 16.5±8 | 18.7±8.5 |
| MDS-UPDRS-III item 11<sup>**,†,‡</sup> |  |  |  |  |
| OFF (/4) | 1.6±1.4 | 0±0<sup>a,b</sup> | 1.6±0.7<sup>a,c</sup> | 2.9±0.9<sup>b,c</sup> |
| ON (/4) | 0.7±0.9 | 0±0<sup>a,b</sup> | 0±0<sup>a</sup> | 1.6±0.6<sup>b</sup> |

Abbreviations: NOFOG, no Freezing of Gait; OFF-FOG, Freezing of Gait in the OFF state only; ONOFF-FOG, Freezing of Gait in the ON and OFF state. <sup>¶</sup>N=43. <sup>\*</sup>,<sup>\*\*</sup>Significant effect of group, <sup>\*</sup>P<0.05; <sup>\*\*</sup>P<0.01. <sup>†</sup>Significant effect of medication state (OFF vs. ON), P<0.01. <sup>‡</sup>Significant group × state interaction, P<0.05. <sup>a,b</sup>Significant difference between listed groups in post-hoc tests, P<0.05. P values are adjusted for PD duration; summary statistics are unadjusted.

### Response of motor scores and change in serum levodopa levels after levodopa challenge in the 3 groups

MDS-UPDRS-III score (item III.11 excluded) varied significantly across medication states (ON vs. OFF) (P<0.01) but did not vary across groups (Table 2). MDS-UPDRS-III total scores calculated including item III.11 varied significantly across states (P<0.01) and groups (P=0.03), with OFF state scores significantly higher in the ONOFF-FOG group compared to the NOFOG group (P=0.03). MDS-UPDRS-III scores decreased by 15.2±7.1 points (range, 5-38) from the “OFF” to “ON” states, corresponding to a percent change of 47±15% (range, 20-80%).

MDS-UPDRS-III item 11 varied significantly across states (P<0.01) and groups (P<0.01), and exhibited a highly significant state × group interaction (P<0.01), as expected by construction.

Among other MDS-UPDRS-III subdomains, significant effects of medication state were identified for all domains except for postural stability, speech, leg agility, and kinetic tremor (Figure 2). Significant effects of group (highest among the ONOFF-FOG group) were identified for Gait (P<0.01), toe tapping (P<0.01), postural stability (P<0.01), speech (P<0.02), leg agility (P<0.03), and pronation/supination of the hands (P<0.05). No statistically-significant state × group interactions were identified for MDS-UPDRS-III items other than FOG. Numerical values of all subdomain scores are summarized in Table S3.

**Figure 2.**
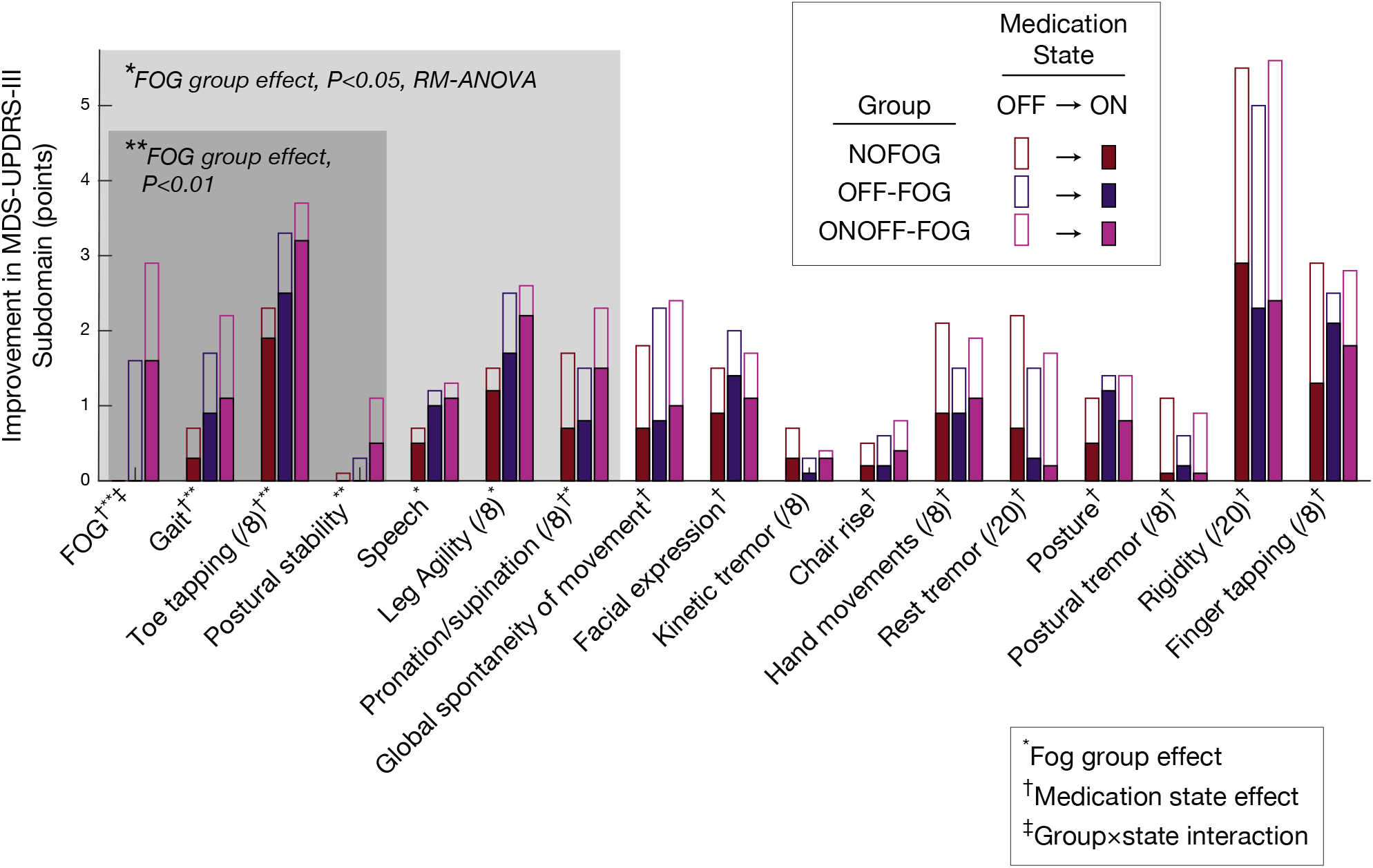
Changes in MDS-UPDRS-III subdomains from OFF to ON medication state in each study group. Empty bars indicate OFF medication scores for each subdomain; superimposed filled bars indicate ON medication scores after levodopa challenge. Study group is indicated by the colors red, blue, and purple for the NOFOG, OFF-FOG, and ONOFF-FOG groups, respectively. Subdomains are sorted in ascending order of group effect P value from left to right. <u>Maximum subdomain scores are 4 points (/4) unless noted</u>. †Significant effect of medication state (OFF vs. ON), P<0.05, RM-ANOVA. *,**Significant effect of group, *P<0.05, **P<0.01. ‡Significant state × group interaction, P<0.05.

Serum levodopa level varied significantly across medication states (“ON” state 27.9 ng/mg vs. “OFF” state 0.3 ng/mg, P<0.01), and in the ON state, was significantly higher in the ONOFF-FOG group compared to the NOFOG group (34.8 vs. 19.9 ng/mg, P=0.01). No other differences were observed between groups.

### Associations between motor features and serum levodopa level in the 3 groups

Linear mixed models revealed a highly significant association between serum levodopa level and MDS-UPDRS-III total score (item 11 excluded; P<0.001; Figure 3) that did not vary across the NOFOG, OFF-FOG, and ONOFF-FOG groups (group effect, P=0.16; group × serum levodopa interaction effect, P=0.32). The estimated slope between MDS-UPDRS-III score and serum levodopa level was −0.52 points•mg/ng (95% CI: −0.62,−0.42). Linear mixed models revealed a highly significant group interaction effect on the association between serum levodopa level with MDS-UPDRS-III item 11 (P<0.001). Estimated slopes between MDS-UPDRS-III item 11 scores and serum levodopa level were 0.00 points•mg/ng (−0.02, 0.02), −0.06 points•mg/ng (−0.07, −0.04), and −0.04 points•mg/ng (−0.05, −0.03) for the NOFOG, OFF-FOG, and ONOFF-FOG groups, respectively. Identified regression parameters and confidence intervals for each group and overall are summarized in Table S4.

**Figure 3.**
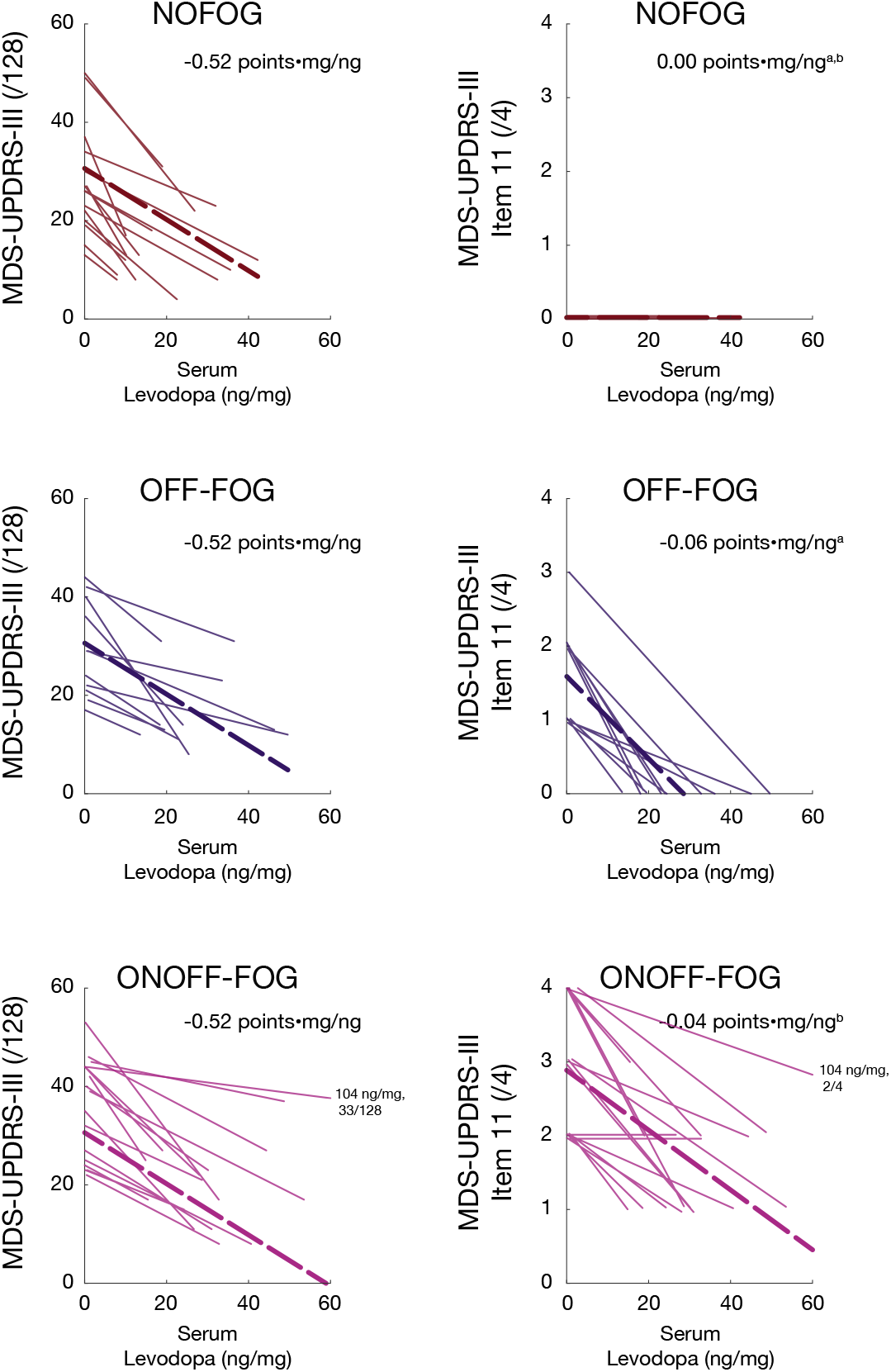
Association between changes in parkinsonian symptom level and changes in serum levodopa level among patients who exhibited a clinically-meaningful response to the acute levodopa challenge. Left: Association between total MDS-UPDRS-III score (excluding item III.11, “Freezing of Gait”) and serum levodopa level. Right: Association between MDS-UPDRS-III item III.11 and serum levodopa level. Study groups are depicted from top to bottom with colors as in Figure 2. Thin lines depict increases in serum levodopa level and corresponding reductions in MDS-UPDRS-III from the OFF to ON states for individual patients. Scores for one patient in the ONOFF-FOG group with a very high ON state serum levodopa level (104 ng/mg) are noted. Bold dashed lines indicate linear regression fits estimated by linear mixed models; numerical values of regression slopes in points•mg/ng calculated by linear mixed models are inset. ^a,b^Significant difference between slope parameters, mixed models, P<0.05. Mixed model results control for PD duration. In the right panel, a small amount of vertical jitter has been added to improve visibility.

## Discussion

In this study we sought to demonstrate that ONOFF-FOG in PD patients is a separate form of FOG, as opposed to OFF-FOG, that is generally not the result of inadequate dopaminergic therapy. We also intended to show that it is not impacted by levodopa in the same manner as other PD motor features. We used rigorous methods that included assessments in the “ON” and “OFF” states. Subjects came to clinic and were assessed in the practically defined “OFF” state. This was followed by a dopaminergic challenge with supratherapeutic medication doses as recommended by others^9^. We used serum levodopa levels as indicators of suprathreshold therapy and required a >20% improvement in MDS-UPDRS III for reaching a true “ON” state. Of the 26 patients in the ONOFF-FOG group 15 reached this threshold. The mean LED given in the levodopa challenge to the ONOFF-FOG group was 451 mg, 28% higher than their standard first dose. Further, 79% of these patients developed dyskinesia. The mean levodopa level was substantially higher in the ONOFF-FOG than the NOFOG and OFF-FOG groups (Table 2). This would strongly suggest that these subjects were adequately treated indicating that the FOG is an “ON” phenomenon and that ONOFF-FOG is an authentic subtype. This subtype is more severe than the OFF-FOG subtype and is associated with more severe impairments to activities of daily living as measured by the MDS-UPDRS-II.

In addition to FOG itself, these results suggest that axial and lower limb parkinsonian signs associated with FOG are also more severe and less responsive to levodopa in ONOFF-FOG, consistent with the interpretation that the overall presentation of ONOFF-FOG may have additional or separate pathophysiology. In addition to FOG retained in the ON state in the ONOFF-FOG group, other related MDS-UPDRS-III subdomains demonstrated limited impact of levodopa, including postural stability, speech, and leg agility, in which the most severe scores in the ON state were in the ONOFF-FOG group. Also, the most significant effects of group were identified for gait, toe tapping, and postural stability, with each being worse in the ONOFF-FOG group. Less significant were leg agility and speech. These findings support the relationship between speech and FOG ^19^. No statistically significant state × group interactions were identified for MDS-UPDRS-III items other than FOG. There were no significant differences in the measures for the cardinal features of tremor, rigidity or bradykinesia of the upper extremities (Table S3). It has been previously shown that FOG does not behave in a similar manner to bradykinesia^20^. This suggests that FOG, sometimes referred to as the fifth cardinal feature^21^, and related features behave differently from other features of PD in relation to levodopa responsiveness. Further, these findings support the notion that FOG, particularly in ONOFF-FOG cases, is governed by a separate pathophysiological mechanism than other cardinal features of PD.

The histories of the individual patients in this study also suggest that ONOFF-FOG can appear without first transitioning through OFF-FOG. The mean duration of FOG in the ONOFF-FOG group was 3.3 years as opposed to 4.4 years for the OFF-FOG group. If ONOFF-FOG was a later result of a cascade of changes then the duration would be expected to be longer. Further, we went back as far as 2006 in record review of the 26 ONOFF-FOG cases to examine if they were first OFF-FOG. Ten subjects had adequate data and clearly showed that they had ONOFF-FOG since FOG inception. This is further support for ONOFF-FOG being a distinct entity as previously suggested^11^. Additional studies with longitudinal follow-up will be necessary to confirm this.

We classified patients by whether or not they had FOG in the “ON” state. We were not attempting to measure gradations of responsiveness or changes in severity. However, it was notable to us that in some of the ONOFF-FOG patients, we saw absolutely no change in FOG severity between the OFF and ON states (n=11), whereas in some we noted what would appear to be improvement (n=15) based on the MDS-UPDRS-III item 11 which is a 0-4 measure of FOG. This caused us to speculate that further subdivisions within the ONOFF-FOG group might be appropriate; however, at this point this remains unclear in part because of the fluctuating nature of the symptoms and in part because item 11 is not validated to measure change. Larger samples and validated measurement scales, perhaps with wearable devices, will be required to investigate this appropriately.

We believe, based on these findings, that research studies of FOG should consider FOG levodopa responsiveness as an important clinical variable. In research projects, subjects are usually divided into those with and without FOG. This common classification approach may be therefore creating admixtures of ONOFF and OFF groups with high intra-group variability, creating conflicting results depending on what subtypes of FOG are more representative. One example of this involves the study of cognitive impairment in FOG. It is generally believed that FOG is associated with executive dysfunction and visuospatial changes and that this association is important from a pathophysiological standpoint^22,23^. However, studies have actually demonstrated that these cognitive measures are associated specifically with ONOFF-FOG^24,25,10,26^ but not OFF-FOG^10, 27^ This inappropriate grouping could also explain why, for example, FOG severity has been associated with reduced functional connectivity within the ‘executive-attention’ neural network in the ‘resting’ state of some studies^28^ but not others^29^. Recent studies have attempted to use cluster analyses to assess subtypes of FOG using imaging, measures of FOG severity, affect and cognition^29, 30^. Perhaps they should include levodopa responsiveness as a variable.

These findings also have important therapeutic implications. FOG observed in patients should be examined carefully to determine if they have OFF-FOG or ONOFF-FOG. One group, OFF-FOG, is treatable, responsive to levodopa and deep brain stimulation^31^. The other, ONOFF-FOG is currently without treatment. A focus on the neurobiology of this particular problem could lead to the development of a useful therapy. From the therapeutic standpoint of a physical or occupational therapist, it is well-established that FOG is a primary contributor to fall risk^32^. Knowledge of whether FOG is levodopa-refractory could redirect therapeutic strategies to reduce fall risk from those focused on medical management to reduce OFF time to those focused on movement strategies to reduce fall risk during unavoidable FOG episodes that could occur at any time.

There are limits to this study. Although this study used rigorous methodology in an attempt to clearly demonstrate the existence of ONOFF-FOG, we did not use blinded raters or randomize “ON” and “OFF” states. Also, FOG tends to be variable in clinical presentation. It would be important to do a test-retest in such subjects to assure consistency of their clinical behavior before and after levodopa challenge. Nevertheless, with careful examination and the use of levodopa blood levels we found consistency of response in the “ON” and “OFF” states and demonstrated the variance of levodopa responsiveness in relation to FOG.

## Conclusions

We believe we have demonstrated the existence of ONOFF-FOG as a distinct subtype of FOG. We also demonstrated that the relationship between FOG and levodopa therapy varies from other cardinal features in PD. Further, ONOFF-FOG is not directly related to OFF-FOG. It would be an important next step to examine the neurobiology of ONOFF-FOG and compare this to other levodopa responsive or induced subtypes.

## Acknowledgements

Motion capture lab assessments were completed by Garrett Alexander M.D., Ph..D..

Serum levodopa levels were completed by the Emory HPLC Bioanalytical Core (EHBC), which was supported by the Department of Pharmacology, Emory University School of Medicine and the Georgia Clinical & Translational Science Alliance of the National Institutes of Health under Award Number UL1TR002378. The content is solely the responsibility of the authors and does not necessarily reflect the official views of the National Institutes of Health

## Supplemental Information

### Levodopa dosages and serum levodopa levels during acute levodopa challenge

Levodopa equivalent dosages used during the acute levodopa challenge are summarized in Table S1. No statistically-significant difference was observed in administered doses between patients who did and who did not subsequently exhibit a clinically-meaningful response (*t*-tests).

Serum levodopa levels after the levodopa challenge are also summarized in Table S1. Serum levodopa levels were significantly higher (increased by ≈79%) among patients who did not exhibit a clinically-meaningful response compared to those who did (P<0.01, *t*-test).

**Table S1.**
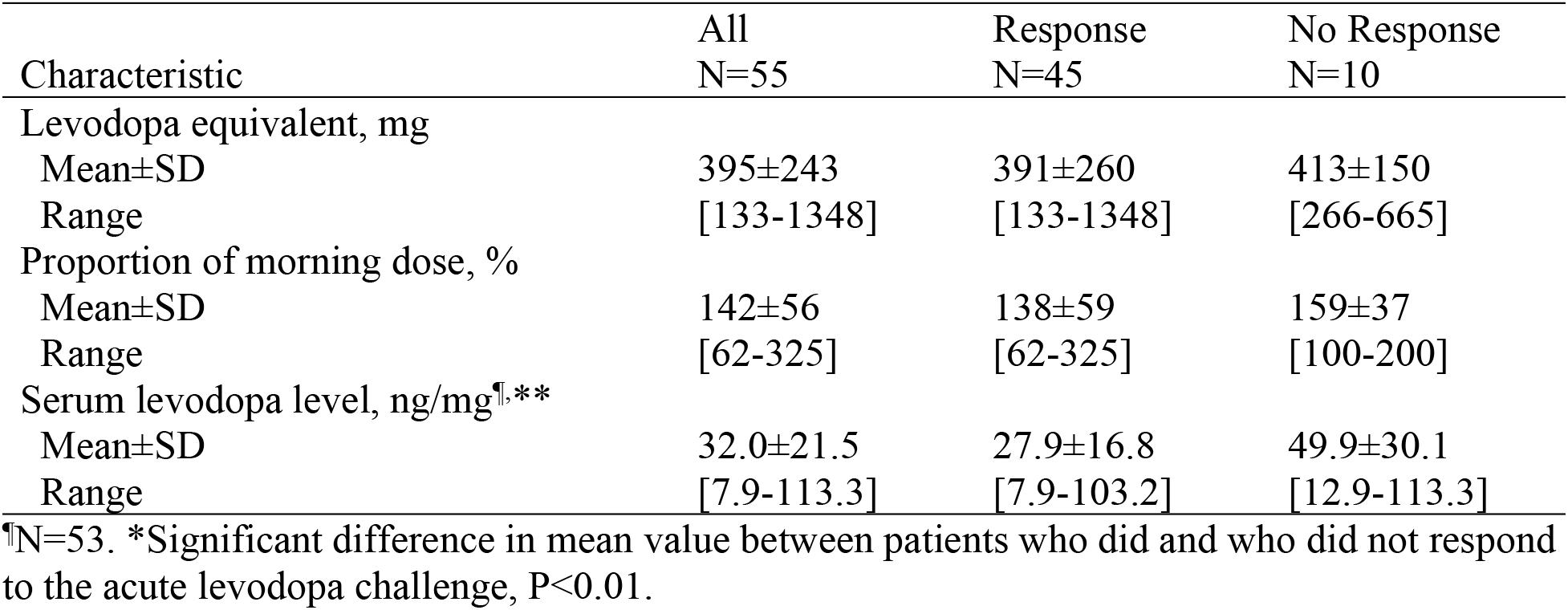
Levodopa equivalent dosages and serum levodopa levels during the levodopa challenge.

| Characteristic | All<br>N=55 | Response<br>N=45 | No Response<br>N=10 |
| --- | --- | --- | --- |
| Levodopa equivalent, mg |  |  |  |
| Mean±SD | 395±243 | 391±260 | 413±150 |
| Range | [133-1348] | [133-1348] | [266-665] |
| Proportion of morning dose, % |  |  |  |
| Mean±SD | 142±56 | 138±59 | 159±37 |
| Range | [62-325] | [62-325] | [100-200] |
| Serum levodopa level, ng/mg<sup>†</sup>,** |  |  |  |
| Mean±SD | 32.0±21.5 | 27.9±16.8 | 49.9±30.1 |
| Range | [7.9-113.3] | [7.9-103.2] | [12.9-113.3] |

<sup>†</sup>N=53. \*Significant difference in mean value between patients who did and who did not respond to the acute levodopa challenge, $P < 0.01$ .

### Demographic and clinical characteristics of all enrolled patients

As described in the main text, all patients were classified into one of three study groups determined *a priori* based on history of the presence of FOG along with scores on the MDS UPDRS-III exam administered before and after the acute levodopa challenge. Analyses reported in the main text consider only those patients who demonstrated an improvement of ≥20% in total MDS-UPDRS-III score after levodopa intake. Demographic and clinical characteristics of all patients initially enrolled are presented here. For simplicity, in this section “ON” refers to assessment after administration of levodopa during the levodopa challenge, and does not necessarily refer to a full “ON” state.

Demographic and clinical characteristics of all patients initially enrolled are summarized in Table S2. Among these, 26 (47%) were in the ONOFF-FOG group, followed by NOFOG 17 (31%), and OFF-FOG 12 (22%). No significant differences between groups were observed in age, sex, education, cognitive ability, age at PD or FOG onset, FOG duration, dyskinesia during the “ON” state (as indicated by the investigator while completing part III of the MDS-UPDRS) or MDS-UPDRS-I or IV. Significant contrasts were observed between the ONOFF-FOG and NOFOG groups on PD duration (10±7 vs. 6±4 y, P<0.05) and levodopa equivalent daily dose (1562±691 vs. 833±302 mg, P<0.01). No significant differences were observed between the OFF-FOG and ONOFF-FOG groups on these variables. The ONOFF-FOG group exhibited significantly more impairment on NFOG-Q (5.5 points, P<0.01) and MDS-UPDRS-II (8.5 points, P<0.01) compared to the OFF-FOG group.

**Table S2.** Demographic and clinical characteristics of all patients initially enrolled, overall and stratified on FOG status.

| Characteristic | All<br>N=55 | NOFOG<br>N=17 | OFF-FOG<br>N=12 | ONOFF-FOG<br>N=26 |
| --- | --- | --- | --- | --- |
| Clinical response |  |  |  |  |
| Yes (n, %) | 45 (82) | 15 (88) | 11 (92) | 19 (73) |
| No (n, %) | 10 (18) | 2 (12) | 1 (8) | 7 (27) |
| Age (y) | 68±9 | 67±12 | 69±5 | 69±8 |
| Sex |  |  |  |  |
| Male (n,%) | 43 (78) | 11 (65) | 10 (83) | 22 (85) |
| Female (n,%) | 12 (22) | 6 (35) | 2 (17) | 4 (15) |
| Education (y) | 16±2 | 17±1 | 18±4 | 16±3 |
| MoCA (/30) | 24.6±4.5<sup>a</sup> | 26.1±3.9 | 25.0±4.0 | 23.5±4.9 |
| PD duration (y)* | 9±6 | 6±4<sup>†</sup> | 11±5 | 10±7<sup>†</sup> |
| Age at PD onset (y) | 59±11 | 61±12 | 58±8 | 59±11 |
| LED (mg)** | 1251±632 | 833±302<sup>†</sup> | 1170±486 | 1562±691<sup>†</sup> |
| MDS-UPDRS-I (/52) | 11.5±5.9<sup>b</sup> | 10.2±4.9 | 11.1±5.5 | 12.9±6.5 |
| MDS-UPDRS-II (/52)** | 16.3±7.3<sup>b</sup> | 11.3±6.7<sup>†</sup> | 16.4±6.2 | 19.8±6.6<sup>†</sup> |
| MDS-UPDRS-IV (/24) | 5.9±3.3<sup>b</sup> | 5.3±3.0 | 6.1±3.3 | 6.3±3.4 |
| ON-state dyskinesia |  |  |  |  |
| Yes (n, %) | 37 (67) | 9 (53) | 11 (92) | 17 (65) |
| No (n, %) | 18 (33) | 8 (47) | 1 (8) | 9 (35) |
| FOG duration (y) | 3±3<sup>c</sup> |  | 4±4 | 3±3 |
| Age at FOG onset (y) | 66±7<sup>c</sup> |  | 65±7 | 66±8 |
| NFOG-Q (/28)** | 20.1±5.1<sup>c</sup> |  | 16.3±5.9<sup>†</sup> | 21.8±3.6<sup>†</sup> |

Abbreviations: NOFOG, no Freezing of Gait; OFF-FOG, levodopa-responsive Freezing of Gait; ONOFF-FOG, levodopa-unresponsive Freezing of Gait. \*Overall effect of group at P<0.05.
\*\*Overall effect of group, P<0.01. <sup>†</sup>Significant difference between listed groups at P<0.05.
<sup>a</sup>N=53. <sup>b</sup>N=54. <sup>c</sup>N=38.

### Response of MDS-UPDRS-III subdomains to levodopa challenge across groups

Mean values of MDS-UPDRS-III subdomains for each medication state in each group are summarized in Table S3. Repeated measures ANOVA models identified statistically significant (P<0.05) effects of medication state on 13/17 MDS-UPDRS-III subdomains investigated. The exceptions (4/17) were postural stability (P=0.07), speech (P=0.14), leg agility (P=0.07), and kinetic tremor (P=0.13).

Significant effects of group were identified for 7/17 MDS-UPDRS-III subdomains investigated: FOG (P<0.01, expected by construction), Gait (P<0.01), toe tapping (P<0.01), postural stability (P<0.01), speech (P<0.02), leg agility (P<0.03), and pronation/supination of the hands (P<0.05). A statistically-significant state × group interaction was identified for MDS-UPDRS-III item 11, FOG; no other significant interactions were identified.

In 6/7 subdomains where significant group effects were identified, subdomain scores increased in severity from NOFOG to OFF-FOG to ONOFF-FOG in both medication states. The only exception was the hand pronation/supination subdomain, in which values among the NOFOG group were slightly higher than those among the OFF-FOG group in the OFF state (1.7±1.3 points vs. 1.5±1.0 points, P=0.43).

**Table S3.** MDS-UPDRS-III items in each medication state, overall and stratified on FOG status among patients who exhibited a clinically-meaningful response to acute levodopa challenge.

| Subdomain | All<br>N=45 | NOFOG<br>N=15 | OFF-FOG<br>N=11 | ONOFF-FOG<br>N=19 |
| --- | --- | --- | --- | --- |
| FOG<sup>†,**,‡</sup> |  |  |  |  |
| OFF (/4) | 1.6±1.4 | 0.0±0.0<sup>a,b</sup> | 1.6±0.7<sup>a,c</sup> | 2.9±0.9<sup>b,c</sup> |
| ON (/4) | 0.7±0.9 | 0.0±0.0<sup>a,b</sup> | 0.0±0.0<sup>a</sup> | 1.6±0.6<sup>b</sup> |
| Gait<sup>†,**,§</sup> |  |  |  |  |
| OFF (/4) | 1.6±0.9 | 0.7±0.7<sup>a,b</sup> | 1.7±0.5<sup>a,c</sup> | 2.2±0.4<sup>b,c</sup> |
| ON (/4) | 0.8±0.7 | 0.3±0.5<sup>b</sup> | 0.9±0.5 | 1.1±0.8<sup>b</sup> |
| Toe tapping<sup>†,**,§</sup> |  |  |  |  |
| OFF (/8) | 3.1±1.3 | 2.3±1.0<sup>b</sup> | 3.3±1.3 | 3.7±1.2<sup>b</sup> |
| ON (/8) | 2.6±1.4 | 1.9±1.2<sup>b</sup> | 2.5±1.3 | 3.2±1.3<sup>b</sup> |
| Postural stability<sup>**</sup> |  |  |  |  |
| OFF (/4) | 0.6±1.1 | 0.1±0.3<sup>b</sup> | 0.3±0.9<sup>c</sup> | 1.1±1.4<sup>b,c</sup> |
| ON (/4) | 0.2±0.7 | 0.0±0.0<sup>b</sup> | 0.0±0.0 | 0.5±1.0<sup>b</sup> |
| Speech<sup>*</sup> |  |  |  |  |
| OFF (/4) | 1.1±0.7 | 0.7±0.6 | 1.2±0.8 | 1.3±0.6 |
| ON (/4) | 0.9±0.7 | 0.5±0.6 | 1.0±0.8 | 1.1±0.7 |
| Leg Agility<sup>*</sup> |  |  |  |  |
| OFF (/8) | 2.2±1.3 | 1.5±0.9<sup>b</sup> | 2.5±1.7 | 2.6±1.2<sup>b</sup> |
| ON (/8) | 1.8±1.4 | 1.2±1.0 | 1.7±1.6 | 2.2±1.4 |
| Pronation/supination<sup>†,*</sup> |  |  |  |  |
| OFF (/8) | 1.9±1.4 | 1.7±1.3 | 1.5±1.0 | 2.3±1.5 |
| ON (/8) | 1.1±1.1 | 0.7±0.8 | 0.8±1.0 | 1.5±1.3 |
| Global spontaneity of movement<sup>†</sup> |  |  |  |  |
| OFF (/4) | 2.2±0.7 | 1.8±0.9 | 2.3±0.5 | 2.4±0.7 |
| ON (/4) | 0.9±0.7 | 0.7±0.5 | 0.8±0.6 | 1.0±0.9 |
| Facial expression<sup>†</sup> |  |  |  |  |
| OFF (/4) | 1.7±0.6 | 1.5±0.9 | 2.0±0.0 | 1.7±0.5 |
| ON (/4) | 1.1±0.7 | 0.9±0.9 | 1.4±0.8 | 1.1±0.6 |
| Kinetic tremor |  |  |  |  |
| OFF (/8) | 0.5±0.9 | 0.7±1.0 | 0.3±0.6 | 0.4±1.0 |
| ON (/8) | 0.2±0.6 | 0.3±0.6 | 0.1±0.3 | 0.3±0.7 |
| Chair rise<sup>†</sup> |  |  |  |  |
| OFF (/4) | 0.7±0.8 | 0.5±0.6 | 0.6±0.7 | 0.8±1.0 |
| ON (/4) | 0.3±0.6 | 0.2±0.6 | 0.2±0.4 | 0.4±0.7 |
| Hand movements<sup>†</sup> |  |  |  |  |
| OFF (/8) | 1.9±1.3 | 2.1±1.3 | 1.5±1.4 | 1.9±1.2 |
| ON (/8) | 1.0±1.0 | 0.9±1.0 | 0.9±1.0 | 1.1±1.0 |
| Rest tremor<sup>†</sup> |  |  |  |  |
| OFF (/20) | 1.8±2.0 | 2.2±2.4 | 1.5±2.2 | 1.7±1.6 |
| ON (/20) | 0.4±0.9 | 0.7±1.2 | 0.3±0.9 | 0.2±0.5 |
| Posture<sup>†</sup> |  |  |  |  |
| OFF (/4) | 1.3±1.0 | 1.1±0.9 | 1.4±1.2 | 1.4±1.0 |
| ON (/4) | 0.8±0.9 | 0.5±0.7 | 1.2±1.2 | 0.8±0.9 |
| Postural tremor<sup>†</sup> |  |  |  |  |
| OFF (/8) | 0.9±1.1 | 1.1±1.2 | 0.6±1.0 | 0.9±1.1 |
| ON (/8) | 0.1±0.4 | 0.1±0.3 | 0.2±0.6 | 0.1±0.2 |
| Rigidity<sup>†</sup> |  |  |  |  |
| OFF (/20) | 5.4±3.6 | 5.5±4.2 | 5.0±3.8 | 5.6±3.2 |
| ON (/20) | 2.5±2.5 | 2.9±2.9 | 2.3±2.9 | 2.4±2.0 |
| Finger tapping<sup>†</sup> |  |  |  |  |
| OFF (/8) | 2.8±1.2 | 2.9±1.5 | 2.5±0.7 | 2.8±1.3 |
| ON (/8) | 1.7±1.2 | 1.3±1.0 | 2.1±0.8 | 1.8±1.4 |

Abbreviations: NOFOG, no Freezing of Gait; OFF-FOG, Freezing of Gait in the OFF state only; ONOFF-FOG, Freezing of Gait in the ON and OFF state. \*\*Significant effect of group, \*P<0.05; \*\*P<0.01. <sup>†</sup>Significant effect of medication state (OFF vs. ON), P<0.05. <sup>‡</sup>Significant group × state interaction, P<0.05. <sup>a,b</sup>Significant difference between listed groups in post-hoc tests, P<0.05. P values are adjusted for PD duration; summary statistics are unadjusted. From top to bottom, subdomains are listed in ascending order of group effect P value.

### Regression parameters identified by linear mixed models

Identified regression parameters and confidence intervals describing associations between MDS-UPDRS-III outcomes and serum levodopa level for each group and overall are summarized in Table S4. Additional linear mixed models applied to MDS-UPDRS-III total scores (item 11 included) revealed a highly significant association between serum levodopa level and MDS-UPDRS-III total score (P<0.001), with a main effect of group (P=0.04) but no evidence of group × serum levodopa interaction (P=0.56).

**Table S4.** Regression parameters describing associations between MDS-UPDRS-III outcomes and serum levodopa level in each study group.

| Outcome/group | Regression parameter |  |
| --- | --- | --- |
|  | Slope (points•mg/ng) | Intercept (points) |
| MDS-UPDRS-III total (item 11 excluded) |  |  |
| All | -0.52 [-0.62, -0.42] | 30.62 [27.62, 33.62] |
| NOFOG | -0.65 [-0.85, -0.46] | 28.00 [22.79, 33.21] |
| OFF-FOG | -0.47 [-0.65, -0.28] | 28.82 [23.11, 34.53] |
| ONOFF-FOG | -0.50 [-0.64, -0.36] | 34.57 [29.73, 39.40] |
| MDS-UPDRS-III item 11 |  |  |
| All | -0.03 [-0.4, -0.02] | 1.52 [1.66, 1.87] |
| NOFOG | -0.00 [-0.02, 0.02]<sup>a,b</sup> | 0.02 [-0.30, 0.34]<sup>a,b</sup> |
| OFF-FOG | -0.06 [-0.07, -0.04]<sup>a</sup> | 1.57 [1.22, 1.92]<sup>a,c</sup> |
| ONOFF-FOG | -0.04 [-0.05, -0.03]<sup>b</sup> | 2.87 [2.57, 3.16]<sup>b,c</sup> |
| MDS-UPDRS-III total |  |  |
| All | -0.57 [-0.67, -0.47] | 32.26 [29.11, 35.41] |
| NOFOG | -0.66 [-0.86, -0.46] | 28.08 [22.77, 33.38]<sup>d</sup> |
| OFF-FOG | -0.53 [-0.72, -0.34] | 30.46 [24.65, 36.27] |
| ONOFF-FOG | -0.54 [-0.69, -0.40] | 37.51 [32.58, 42.43]<sup>d</sup> |

Regression parameters were determined from mixed models and control for PD duration. Regression parameters are presented as estimate [95% confidence interval]. <sup>a-d</sup>Significant contrast between marked groups, $P<0.05$ .

